# Bridging LLM Reasoning and Chemical Knowledge via an Evolutionary Multi-Agent Framework for Molecular Synthesis

**DOI:** 10.64898/2026.05.02.722342

**Authors:** Yicong Chen, Jiahua Rao, Jiancong Xie, Youhan Sun, Yuedong Yang

**Affiliations:** School of Computer Science and Engineering, Sun Yat-sen University, 510000, Guangzhou, China; Guangdong Provincial Key Laboratory of Computational Science, Sun Yat-sen University, 510000, Guangzhou, China

## Abstract

**Motivation:** Molecular design faces the dual challenge of navigating a vast chemical space while ensuring experimental synthesizability. Traditional models are constrained by small datasets, restricting their scalability and broader chemical context. In contrast, Large Language Models (LLMs) encapsulate extensive synthesis protocols derived from vast scientific literature, yet they struggle to leverage this potential due to severe hallucinations and a superficial grasp of rigorous chemical logic.

**Results:** We propose EvoSyn, an evolutionary multi-agent framework that synergizes LLM reasoning with domain experts for preference-aware molecular synthesis. EvoSyn orchestrates a dual-process evolutionary paradigm: a co-evolving process that collaboratively aligns linguistic capabilities with multi-objective constraints, and a self-evolving process formulated as a Markov Game. Through evolution and reinforcement learning, agents actively learn from mistakes, utilizing domain feedback to penalize invalid proposals and ground generation in feasible reaction pathways. Extensive evaluations on comprehensive benchmarks demonstrate that EvoSyn significantly outperforms state-of-the-art baselines. These results highlight that by integrating LLM-guided self-evolution with rigorous domain validation to mitigate hallucinations, EvoSyn effectively yields molecules that are both bioactive and synthetically actionable.

**Availability and implementation:** Implementation code is available as supplementary material.

**Supplementary information:** Supplementary data are available at Bioinformatics online.

## 1. Introduction

Navigating the immense chemical space, estimated to exceed 10^60^ pharmacologically active molecules Bohacek et al. (1996), has long been a fundamental challenge in drug discovery. While recent advances in Deep Learning (DL) have empowered generative models to rapidly explore this vast landscape, a substantial chasm remains between theoretical design and experimental feasibility Sanchez-Lengeling and Aspuru-Guzik (2018). The critical bottleneck is that generated molecules often lack practical synthesizability guarantees. This deficiency severely restricts their real-world application, leaving high-potential candidates trapped *in silico*, unreachable by current laboratory synthesis methods.

To bridge the gap between generation and synthesis, existing strategies generally fall into three categories: (1) integrating heuristic or learned scoring functions (e.g., SAscore, DeepSA Ertl and Schuffenhauer (2009); Coley et al. (2018)) into optimization objectives; (2) employing forward synthesis models (e.g., SynNet, Synformer Gao et al. (2021, 2025)) that assemble molecules from building blocks via predefined templates; and (3) leveraging Large Language Models (LLMs) (e.g., SynLlama Sun et al. (2025)) for chemical text processing. However, traditional domain models are typically constrained by small datasets, restricting their scalability and broader chemical context.

While LLMs circumvent these data constraints, their deployment for robust synthesis planning faces significant hurdles arising from a superficial utilization of chemical knowledge. Relying predominantly on statistical pattern matching rather than explicit reaction logic, LLMs frequently suffer from hallucination, generating plausible-appearing but experimentally unfeasible structures Guo et al. (2023). Although integrating external expert tools offers potential mitigation Bran et al. (2023); Boiko et al. (2023), achieving seamless synergy remains elusive. Current paradigms typically lack mechanisms for deep collaboration, as the LLM and expert tools often operate in decoupled environments without a shared feedback loop to correct errors dynamically Edwards et al. (2022).

Furthermore, the absence of evolutionary mechanisms prevents systems from “learning from mistakes” via continuous interaction. Most frameworks remain static post-training, unable to accumulate experience or adaptively optimize reasoning under complex, multi-objective constraints Shinn et al. (2023); Madaan et al. (2023). This rigidity hinders alignment with dynamic user preferences, limiting effective long-term navigation of the intricate chemical space.

To address these limitations, we introduce **EvoSyn**, an evolutionary multi-agent framework designed for preference-aware molecular design and synthesis planning. Unlike static approaches, EvoSyn orchestrates four specialized agents (preference interpreter, synthesis planner, reasoning responder, and consistency evaluator) within a dual-process evolutionary paradigm to bridge the gap between LLM-based reasoning and rigorous chemical validation. In the co-evolving process, agents leverage a feedback loop to correct errors and balance synthesis preferences in real time, thereby transforming the static capabilities of large language models into dynamic and expert-level proficiency. In the self-evolving process, molecular synthesis is formulated as a Markov Game Littman (1994), wherein synthesis planner agent govern the game-theoretic training among sub-agents. Through continuous optimization under specific multi-objective constraints, EvoSyn ultimately evolves a strong preference-aware capability for molecular synthesis.

Extensive evaluations across reconstruction benchmarks, preference-aware planning, and synthesis optimization tasks demonstrate that EvoSyn consistently outperforms state-of-the-art (SOTA) baselines, generating molecules that are not only bioactive and structurally valid but also synthetically actionable by design. Our contributions are summarized as:

- We propose **EvoSyn**, an evolutionary framework that bridges linguistic reasoning with chemical validation under the guidance of LLMs, enabling dynamic trade-offs for preference-aware synthesis.
- We formulate molecular synthesis as a Markov Game, empowering agents to perform adaptive multi-objective optimization and self-evolve to generate rigorous, experimentally actionable synthetic routes.
- We validate EvoSyn through comprehensive experiments evaluating both synthetic route planning ability and preference-conditioned molecular synthesis, demonstrating that it achieves SOTA performance across all metrics.

## 2. Related works

### 2.1. Molecular synthesis

The pursuit of generating synthetically feasible molecules has driven the development of diverse strategies. Optimization-based methods integrate heuristic or learned scoring functions (e.g., SAscore, DeepSA Ertl and Schuffenhauer (2009); Coley et al. (2018)) into the objectives of generative models Gómez-Bombarelli et al. (2018); Hoogeboom et al. (2022) to guide the search toward simpler structures. Forward synthesis models, such as SynNet and Synformer Gao et al. (2021, 2025), enforce synthesizability by constructing molecules from building blocks via predefined reaction templates. More recently, Language-based approaches (e.g., MolT5, SynLlama Edwards et al. (2022); Sun et al. (2025)) treat synthesis planning as a text generation task, leveraging the linguistic capabilities of LLMs. However, these methods remain constrained by a superficial utilization of chemical knowledge. They predominantly learn statistical correlations from training corpora rather than internalizing underlying reaction mechanisms. It is precisely this deficiency that we address by integrating explicit chemical reasoning and evolutionary reinforcement learning, enabling the system to ensure rigorous synthetic validity.

### 2.2. Multi-agent systems

The deployment of LLMs is shifting from static interaction to dynamic, agent-based collaboration. Multi-agent frameworks (e.g., CAMEL Li et al. (2023)) have demonstrated that simulating human-like role-playing significantly enhances problem-solving in complex tasks. In the scientific domain, Coscientist Boiko et al. (2023) and ChemCrow Bran et al. (2023) have pioneered the integration of LLMs with domain-specific tools. More recent approaches, such as SciAgents Ghafarollahi and Buehler (2025) and ChemMAS Yang et al. (2025b), further explore role-based collaboration to decompose complex reasoning tasks into sub-routines handled by specialized agents. Despite these advancements, current paradigms face significant limitations in molecular design. Most chemical agents, such as MT-MOL Kim et al. (2025), operate as static “tool users” with decoupled interactions, lacking mechanisms for deep collaboration or dynamic strategy refinement. Furthermore, existing self-correction methods typically rely on prompt engineering or simple iterative feedback rather than game-theoretic reinforcement learning Shinn et al. (2023). This prevents them from effectively “learning from mistakes” to balance conflicting objectives (e.g., bioactivity vs. synthesizability) in a rigorous chemical context.

## 3. Materials and methods

### 3.1. The EvoSyn framework

We formulate preference-aware molecular synthesis as a conditional generation task. Given a natural language query *q*, which encodes both explicit constraints and implicit preferences, and an optional reference molecule *m*_ref_, the goal is to generate an optimal synthesis plan. The output comprises a synthetic route *τ* (a sequence of reaction steps) and a reasoning response *o* (theoretical justification). Formally, the objective is to learn a policy *π* that maximizes a joint reward function *R*(*τ, o* | *q, m*_ref_), reflecting the alignment between the generated output and multi-objective preferences.

Directly optimizing *R* in a single step is intractable due to the combinatorial complexity of the reaction space. Therefore, we decompose the generation and optimization process into a dual-process evolutionary paradigm:

- **Co-Evolving Process:** Multiple agents collaboratively perform iterative multi-objective trade-offs to refine the specific route *τ* for the current query, aligning broad linguistic knowledge with specific constraints.
- **Self-Evolving Process:** We formulate the task as a Markov Game, utilizing reinforcement learning to drive the self-evolution of agent. This updates the policy *π* by learning to achieve global optimality.

To execute this paradigm, EvoSyn orchestrates four specialized agents, as illustrated in Figure 1.a:

1. **Preference Interpreter** *ϕ*_intp_ : Acts as the intent parser, extracting explicit constraints and implicit preferences from query *q* to guide the synthesis direction.
2. **Synthesis Planner** *ψ*_plan_ : Serves as the core generative engine, leveraging the LLM to propose concrete synthetic routes and reaction steps.
3. **Reasoning Responder** *ρ*_res_ : Provides explicit chemical reasoning and theoretical justifications (*o*) for the proposed steps, bridging generation and understanding.
4. **Consistency Evaluator** *ξ*_eval_ : as the rigorous critic by connecting with external validation tools to quantitatively assess the generated routes against objectives.

**Figure 1.**
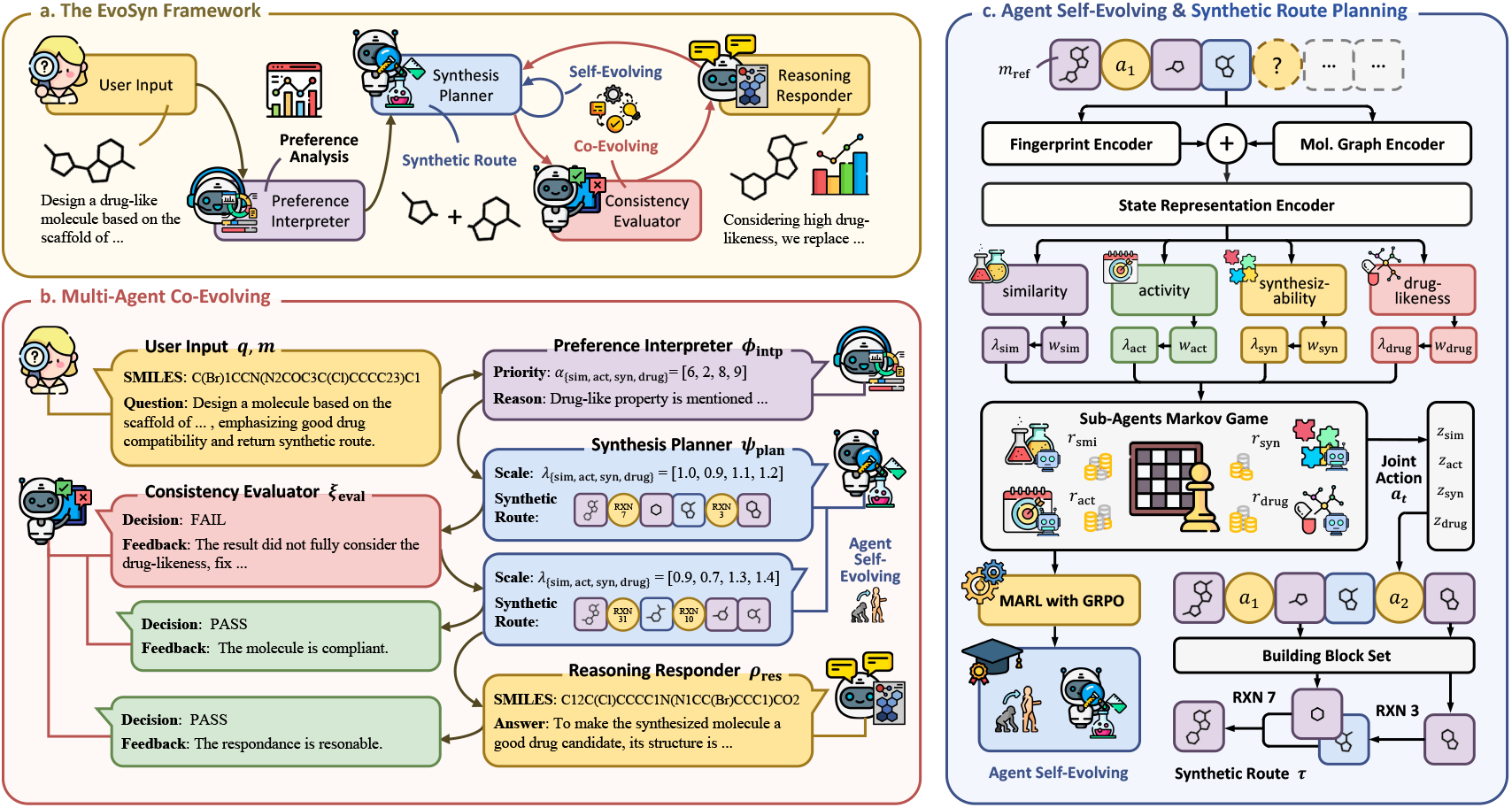
Overview of EvoSyn. (a) the overall framework architecture, (b) the co-evolutionary interaction between multi-agents, and (c) the agent self-evolution mechanism for synthetic route planning.

### 3.2. Collaborative evolution via iterative refinement

To efficiently execute molecular synthesis, the agents in EvoSyn first infer the synthesis preferences embedded in the semantic information of the input, and then dynamically balance multiple potentially conflicting objectives, continuously refining the synthetic route and outputs. Instead of a static generation process, this mechanism orchestrates the agents to continuously refine the synthetic route and outputs via Synthesis Initialization and Iterative Refinement, as shown in Figure 1.b.

#### 3.2.1. Synthesis initialization

In this stage, the preference interpreter *ϕ*_intp_ takes the input query *q* and maps it to a preference representation:

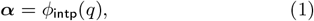

where ***α*** ∈ ℝ^|*N*|^ is the priority vector, and |*N*| denotes the number of predefined synthesis objectives.

Conditioned on ***α***, the synthesis planner *ψ*_plan_ produces a scale vector ***λ***_0_ to modulate the objective weights of sub-agents, subsequently sampling an initial synthetic route *τ*_0_:

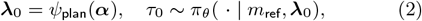

where *π*_*θ*_ denotes the joint policy distribution of the sub-agents in *ψ*_plan_ (detailed in Section 3.3.1).

Finally, the reasoning responder *ρ*_res_ generates the answer *o*_0_ by maximizing the conditional likelihood:

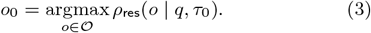

However, relying solely on this one-shot initialization often yields suboptimal solutions that fail to balance conflicting objectives (e.g., high bioactivity vs. low synthesis cost). Consequently, the system proceeds to the iterative refinement phase to further enhance the synthesis plan.

#### 3.2.2. Iterative refinement

In this stage, the system enters an iterative loop to address complex trade-offs dynamically. In the *k*-th iteration of co-evolving, the consistency evaluator *ξ*_eval_ generates feedback *δ*_*k*_ based on query *q* and synthetic route *τ*_*k*_:

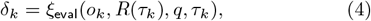

where *R*(*τ*_*k*_) denotes the application of the reward function *R*(·) to quantify the performance of *τ*_*k*_ with respect to different objectives.

Given the feedback *δ*_*k*_, the synthesis planner *ψ*_plan_ and the reasoning responder *ρ*_res_ collaboratively adjust their strategies to produce an improved route *τ*_*k*+1_ and output *o*_*k*+1_:

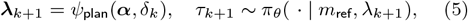

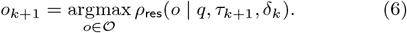

This process repeats until the consistency evaluator determines that the synthetic route satisfies the requirements or a maximum iteration depth is reached.

### 3.3. Self-evolution via synthesis markov game

Despite the effectiveness of collaborative evolution in navigating trade-offs, the system remains reliant on the base LLM, which is susceptible to hallucinations and lacks the deep domain-specific optimization required for rigorous synthesis planning. To overcome these limitations, we formulatethe multi-objective molecular synthesis task as a Markov Game (MG), where reinforcement learning drives the *self-evolution* of the agents. Specifically, Group Relative Policy Optimization (GRPO) Shao et al. (2024) is employed to optimize the policy of each sub-agent in the synthesis planner. Through this formulation, the synthesis planner converges to a policy that is jointly optimal across multiple objectives, thereby enabling robust self-evolving behavior.

#### 3.3.1. Markov game for molecular synthesis

As illustrated in Figure 1.c., the input molecule *m*_ref_ serves as the initial state *m*_0_. At each subsequent time step *t*, the agent selects a reaction template conditioned on the current molecular state *m*_*t*_, applying the corresponding inverse reaction to decompose the molecule into simpler precursors *m*_*t*+1_. This decomposition process repeats iteratively until all resulting fragments belong to a predefined set of commercially available building blocks. The resulting trajectory is then reversed to yield the final forward synthetic route *τ*.

Since *m*_*t*+1_ depends only on the previous state *m*_*t*_, the process satisfies the Markov property and can be naturally modeled as a Markov Game (MG). The MG is characterized by the following key components:

- **Agents**: The sub-agent set is 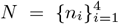. Each sub-agent *n*_*i*_ focuses on a specific property of the synthesized molecule, including similarity, activity, synthesizability, and drug-likeness.
- **State**: The state *s*_*t*_ ∈ *S* represents the current molecule *m*_*t*_ along the synthetic route.
- **Action**: The joint action *a*_*t*_ represents the selection of a reaction template. *a*_*t*_ is sampled from the joint action space *A* according to the joint policy distribution *π*_*θ*_(*a*_*t*_|*s*_*t*_), i.e., *a*_*t*_ ∼ *π*_*θ*_(· | *s*_*t*_). This joint distribution is formulated as:

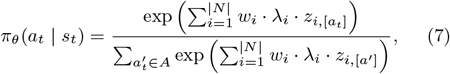

where 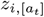 denotes the logit value produced by sub-agent *n*_*i*_ for action *a*_*t*_.

- **Transition**: The transition probability function *P* : *S × A* → *S* is governed by chemical reaction rules, meaning the molecule *m*_*t*_ is transformed to *m*_*t*+1_ through a reaction.
- **Reward**: The joint reward function is defined as *R* : *S × A× S* → ℝ^|*N*|^. Each reward *r*_*i*_ ∈ *R* is assigned to the sub-agent *n*_*i*_ that focuses on the corresponding property. Specifically:

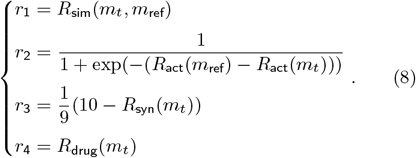

The details of reward functions are shown in Supplementary Section S1.2.

#### 3.3.2. Policy model architecture

To balance parameter efficiency with objective-specific specialization, the policy model leverages a shared-backbone design. For the set of sub-agents *N*, the policy models 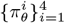 share a common Molecular Encoder and State Representation Encoder to extract unified semantic features, while employing independent Policy Heads to generate distinct action distributions aligned with their sub-goals.

##### Molecular fingerprint encoder

Each molecule is represented using a 1,024-bit Extended Connectivity Fingerprint (ECFP) with a radius of 2 Rogers and Hahn (2010). These fingerprints are processed by a Multi-Layer Perceptron (MLP) to extract molecular hidden features *h*_fp_:

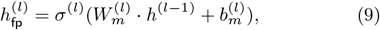

where *l* is the *l*-th layer of MLP, *σ* is the activation function, and *W*_*m*_ and *b*_*m*_ are learnable parameters of the MLP.

##### Molecular graph encoder

For molecules represented as graph structures, a Graph Isomorphism Network (GIN) Xu et al. (2018) is implemented. The GIN updates the features of each atom *e*_*v*_ in the molecule *m* and an add pooling operation aggregates the features into *h*_graph_:

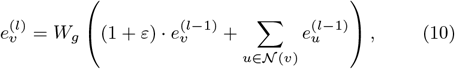

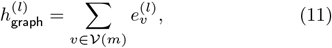

where *W*_*g*_ is a learnable parameter, *ε* is a fixed scaling factor, 𝒩 (*v*) denotes the neighbors of atom *v* and 𝒱 (*m*) denotes the set of all atoms in molecule *m*.

##### State representation encoder

Given the last layer molecular hidden representations *h*_fp_ and *h*_graph_, we integrate them to construct a unified feature representation *x*:

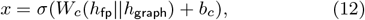

where *W*_*c*_ and *b*_*c*_ denote learnable parameters.

##### Multi-objective policy heads

For each sub-agent *n*_*i*_, an agent-specific prediction head is employed to compute the logit *z*_*i*_ for molecule *m* under the corresponding target:

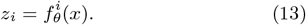

The resulting logit *z*_*i*_ is subsequently aggregated into the joint logit following the procedure described in Section 3.3.1, from which the next joint action *a* is sampled.

#### 3.3.3. GRPO-based self-evolutionary training

We employ Group Relative Policy Optimization (GRPO) Schulman et al. (2017) to optimize its policy model 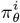 effectively. For each sub-agent *n*_*i*_, the GRPO loss is defined as:

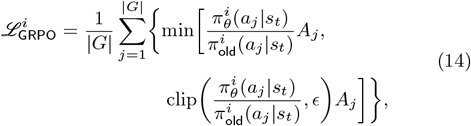

where 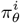 denotes the policy model of sub-agent *n*_*i*_, clip(·, *ϵ*) is a clipping operator with ratio *ϵ* used to stabilize training.

The advantage *A*_*j*_ and the KL divergence term 𝔻_KL_[·] are:

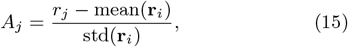

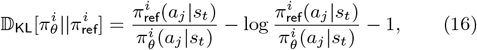

where **r**_*i*_ denotes the reward vector over the sample set *G*. The reward **r**_*i*_ is determined by the specific property focused on by sub-agent *n*_*i*_.

The parameters *θ* are updated to maximize the advantages across all sub-agents. The evolutionary loss function is:

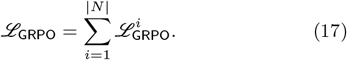

By minimizing this loss, the shared backbone evolves to capture the common chemical reasoning patterns, while the policy heads learn to satisfy the diverse multi-objective constraints defined by the sub-agents.

## 4. Results

### 4.1. Experimental setup

#### Reaction templates and building blocks

We follow the setting of Gao et al. (2025), employing 115 reaction templates that cover common uni-, bi-, and tri-molecular reactions, together with 223,244 building blocks from Enamine’s U.S. stock catalog. More details are shown in Supplementary Section S1.3.

#### Datasets

We evaluate the synthesis capability of EvoSyn on widely used datasets: Enamine REAL ena and ChEMBL Gaulton et al. (2012). Following Gao et al. (2025), the evaluation is conducted on 1,000 molecules randomly sampled from each dataset, respectively. The details are discussed in Supplementary Section S2.5.

We introduce SynQA, a question-answering benchmark for chemical synthesis consisting of two subsets. SynQA-Basics contains 986 QA pairs focusing on mechanistic knowledge of chemical synthesis. SynQA-Design comprises 240 carefully curated, high-quality preference-conditioned molecular synthesis questions, designed to assess the ability to balance multiple molecular properties in multi-objective synthesis scenarios. More details about SynQA are shown in Supplementary Section S2.6.

#### Baselines

In synthesizable molecule reconstruction benchmarks, we evaluate EvoSyn against state-of-the-art molecular synthesis and retrosynthesis baselines, including ReaSyn Lee et al. (2025), SynFormer Gao et al. (2025), SynLlama Sun et al. (2025), ChemProjector Luo et al., SynNet Gao et al. (2021), and PDVN Liu et al. (2023). On the SynQA dataset, we adopt Qwen3-235B-A22B Yang et al. (2025a), GPT-5.2 Singh et al. (2025), Gemini3-Pro dee, and Deepseek-R1 Guo et al. (2025) as large language model baselines under the zero-shot setting. On SynQA-Design, we fine-tune two representative models on a smaller scale, Qwen2.5-7B (FT) and Llama3.1-8B (FT), which serve as fine-tuned baselines for comparison. In addition, we evaluate the performance of the state-of-the-art chemical multi-agent baselines ChatDrug Liu et al. (2024) and ChemCrow Bran et al. (2023).

#### Metrics

Reconstruction Rate and Similarity are utilized to evaluate synthesizable molecule reconstruction performance. The Reconstruction Rate represents the proportion of successfully re-synthesized molecules, while Similarity quantifies the resemblance between the synthesized and original molecules. Accuracy and F1 Score (*F*_1_) serve as the evaluation metrics for SynQA-Basics. Given the absence of ground-truth answers in SynQA-Design, we employ Mean Rank (MR) and Mean Reciprocal Rank (MRR) to assess the relative performance of different methods. The Vina Score Eberhardt et al. (2021) is used to measure the binding affinity of optimized molecules against specific targets.

### 4.2. Evaluation on molecule reconstruction

To validate the fundamental synthesizability ensured by the self-evolutionary mechanism, we first evaluate EvoSyn on standard reconstruction benchmarks, following the protocols of Gao et al. (2025) and Lee et al. (2025).

As shown in Table 1, EvoSyn consistently outperforms all molecular synthesis and retrosynthesis baseline methods, yielding substantial improvements across all benchmarks. Specifically, EvoSyn achieves the highest Reconstruction Rates of 83.7% and 28.8% on Enamine REAL and ChEMBL, respectively, surpassing the previous state-of-the-art by 21.1% and 39.1%. Even in cases where reconstruction fails, EvoSyn maintains the highest Similarity scores, reaching 0.9597 and 0.6984 on the three datasets. Notably, the overall performance on ChEMBL is relatively low for all methods. This may be attributed to the presence of molecules beyond the predefined synthesizable space (i.e., requiring reactions or building blocks not provided in the reference set) Lee et al. (2025). Nevertheless, EvoSyn still demonstrates clear superiority over all baseline methods.

**Table 1.**
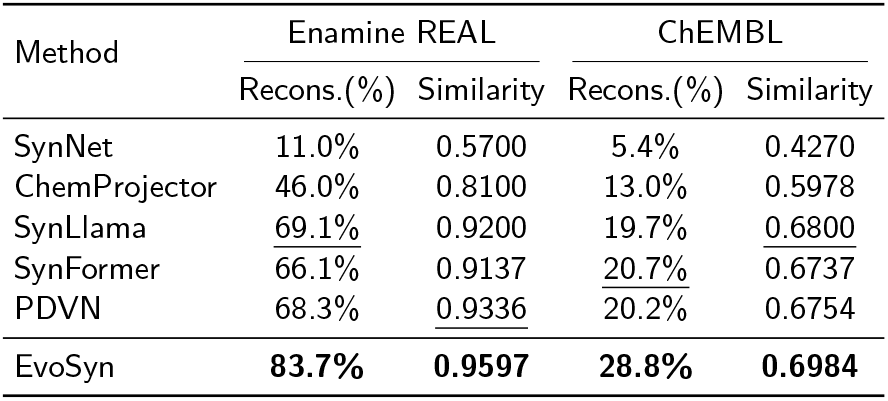
Reconstruction Rate (Recons.) and Similarity on Enamine REAL and ChEMBL.

| Method | Enamine REAL |  | ChEMBL |  |
| --- | --- | --- | --- | --- |
|  | Recons.(%) | Similarity | Recons.(%) | Similarity |
| SynNet | 11.0% | 0.5700 | 5.4% | 0.4270 |
| ChemProjector | 46.0% | 0.8100 | 13.0% | 0.5978 |
| SynLlama | 69.1% | 0.9200 | 19.7% | 0.6800 |
| SynFormer | 66.1% | 0.9137 | 20.7% | 0.6737 |
| PDVN | 68.3% | 0.9336 | 20.2% | 0.6754 |
| EvoSyn | **83.7%** | **0.9597** | **28.8%** | **0.6984** |

These results corroborate that the self-evolution driven by reinforcement learning effectively enables EvoSyn to internalize rigorous chemical rules, providing a robust structural foundation for downstream synthesis planning.

### 4.3. Evaluation on preference-aware synthesis planning

We utilize the SynQA dataset to assess the model’s comprehension of reaction mechanisms and its capability to perform preference-aware synthesis under complex constraints.

To provide a comprehensive benchmarking, we compare EvoSyn and EvoSyn (FT) against zero-shot generalist LLMs, fine-tuned specific LLMs and chemical multi-agent methods. As illustrated in Table 2 and Table 3, EvoSyn and EvoSyn (FT) achieve the best performance on both subsets among all methods. On SynQA-Basics, EvoSyn attains an Accuracy of 0.8469 and an *F*_1_ score of 0.8394 under the zero-shot setting, outperforming the zero-shot LLM baselines and fine-tuned 7B/8B LLMs. EvoSyn (FT) further improved the Accuracy and *F*_1_ score to 0.8979 and 0.8935 through fine-tuning. On SynQA-Design, which involves open-ended planning, EvoSyn achieves an MR of 1.6327 and an MRR of 0.7631, representing improvements of 15.6% and 19.6% over the strongest baseline method.

**Table 2.** Accuracy and *F*_1_ on SynQA-Basics.

| Model | SynQA-Basics |  |
| --- | --- | --- |
| | Accuracy ↑ | $F_1$ ↑ |
| Qwen3-235B-A22B | 0.6775 ± 0.0322 | 0.6779 ± 0.0316 |
| Gemini-3-Pro | 0.7346 ± 0.0146 | 0.7335 ± 0.0135 |
| GPT-5 | 0.6428 ± 0.0234 | 0.6428 ± 0.0291 |
| DeepSeek-R1 | 0.7142 ± 0.0285 | 0.7156 ± 0.0218 |
| Qwen2.5-7B (FT) | 0.7669 ± 0.0107 | 0.7622 ± 0.0103 |
| LLaMA-3.1-8B (FT) | 0.7573 ± 0.0073 | 0.7582 ± 0.0097 |
| EvoSyn | 0.8469 ± 0.0134 | 0.8394 ± 0.0139 |
| EvoSyn (FT) | **0.8979 ± 0.0115** | **0.8935 ± 0.0126** |

**Table 3.**
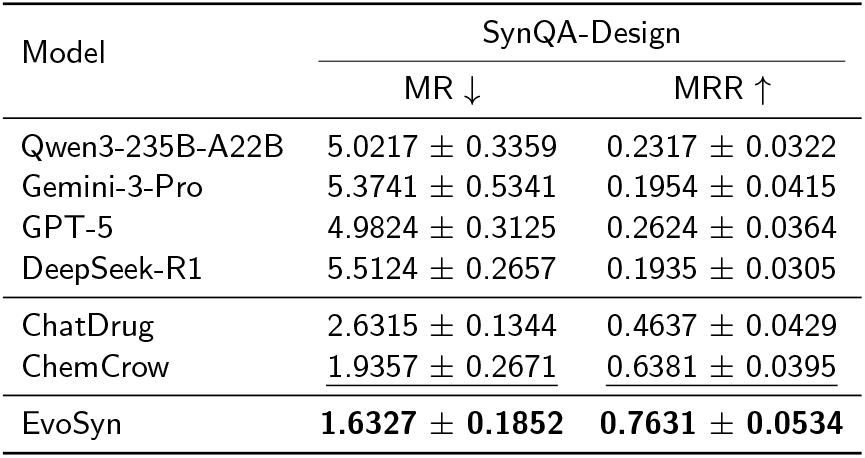
Mean Rank (MR) and Mean Reciprocal Rank (MRR) on SynQA-Design.

| Model | SynQA-Design |  |
| --- | --- | --- |
| | MR ↓ | MRR ↑ |
| Qwen3-235B-A22B | 5.0217 ± 0.3359 | 0.2317 ± 0.0322 |
| Gemini-3-Pro | 5.3741 ± 0.5341 | 0.1954 ± 0.0415 |
| GPT-5 | 4.9824 ± 0.3125 | 0.2624 ± 0.0364 |
| DeepSeek-R1 | 5.5124 ± 0.2657 | 0.1935 ± 0.0305 |
| ChatDrug | 2.6315 ± 0.1344 | 0.4637 ± 0.0429 |
| ChemCrow | 1.9357 ± 0.2671 | 0.6381 ± 0.0395 |
| EvoSyn | **1.6327 ± 0.1852** | **0.7631 ± 0.0534** |

These experiments demonstrate that through multi-agent co-evolution, EvoSyn effectively captures the nuances of reaction mechanisms and exhibits strong reasoning capabilities in aligning synthesis plans with implicit user preferences.

### 4.4. Evaluation on lead synthetic accessibility optimization

For further validating the synthesizable optimization capacity of EvoSyn, we employ EvoSyn to optimize the initial lead candidates generated by Pocket2Mol Peng et al. (2022) for 15 targets within the LIT-PCBA dataset, following the experimental protocol of Luo et al.. Table 4 summarizes the Vina Score (binding affinity) and Similarity metrics.

**Table 4.** Vina Score and Similarity on LIT-PCBA.

| Target | Vina Score (kcal/mol) |  |  | Similarity |  |
| --- | --- | --- | --- | --- | --- |
|  | Pocket-2Mol | Chem-Projector | EvoSyn | Chem-Projector | EvoSyn |
| ADRB2 | -8.70 | -9.09 | **-11.09** | 0.5804 | **0.5914** |
| ALDH1 | -9.10 | -9.00 | **-11.28** | 0.3875 | **0.4619** |
| ESR1_ago | -9.46 | -9.36 | **-12.78** | 0.4066 | **0.4637** |
| ESR1_ant | -9.77 | -9.77 | **-11.88** | 0.4965 | **0.5915** |
| FEN1 | -6.54 | -6.60 | **-8.20** | 0.4180 | **0.4205** |
| GBA | -6.96 | -7.69 | **-9.55** | 0.3140 | **0.4505** |
| IDH1 | -9.50 | -9.56 | **-12.68** | 0.3830 | **0.5143** |
| KAT2A | -8.07 | -8.18 | **-10.37** | 0.5028 | **0.5767** |
| MAPK1 | -8.61 | -8.98 | **-11.03** | 0.4274 | **0.4293** |
| MTORC1 | -10.17 | -10.20 | **-11.77** | 0.4565 | **0.4633** |
| OPRK1 | -8.36 | -8.77 | **-10.70** | 0.5125 | **0.5829** |
| PKM2 | -9.45 | -10.09 | **-12.43** | 0.4558 | **0.4668** |
| PPARG | -8.47 | -8.53 | **-10.11** | 0.4992 | **0.5321** |
| TP53 | -7.11 | -7.47 | **-8.13** | 0.5566 | **0.5688** |
| VDR | -10.51 | -10.46 | **-13.02** | 0.4355 | **0.4805** |
| Average | -8.72 | -8.92 | **-11.00** | 0.4555 | **0.5063** |

EvoSyn successfully converts all molecules generated by Pocket2Mol into synthetically accessible analogs, thereby making the synthesis of the lead molecules feasible. Additionally, the synthetically accessible analogs consistently achieve superior Vina scores compared to those generated by Pocket2Mol and optimized by ChemProjector. On average, EvoSyn reduces the Vina Score by 2.08 relative to the baseline. The largest improvement is observed on target *ESR1 ago*, where the molecules optimized by EvoSyn reduce the Vina Score by 3.42, corresponding to a 36.5% increase in activity. While optimizing synthetic accessibility and enhancing activity, EvoSyn maintains the highest molecular similarity among all methods, thereby preserving the structural integrity of the lead compounds.

These results highlight that our evolutionary multi-agent system effectively navigates the critical trade-off among synthetic accessibility, bioactivity enhancement and structural preservation, validating the efficacy of the multi-objective optimization inherently modeled in our Markov Game formulation.

### 4.5. Ablation study

To explicitly quantify the contribution of each component in our framework, we conduct ablation studies focusing on three key aspects: the effectiveness of the co-evolving agents, the impact of self-evolving with fine-tuning, and the contribution of self-evolving with reinforcement learning.

#### Contribution of co-evolving agents

We first investigate the necessity of the multi-agent collaboration mechanism, with results detailed in Figure 2 a1 and Supplementary Table S2. The Synthesis Planner *ψ*_plan_ emerges as the most critical component. Specifically, its removal precipitates a sharp decline in Accuracy by 22.4% compared to the full model. This substantial drop underscores that the planner is not merely an initialization step, it integrates deep domain-specific knowledge and synthetic constraints that guide the subsequent generation. Without this structured reasoning path, the system struggles to balance conflicting objectives from the outset, leading to chemically infeasible or suboptimal synthetic routes.

**Figure 2.**
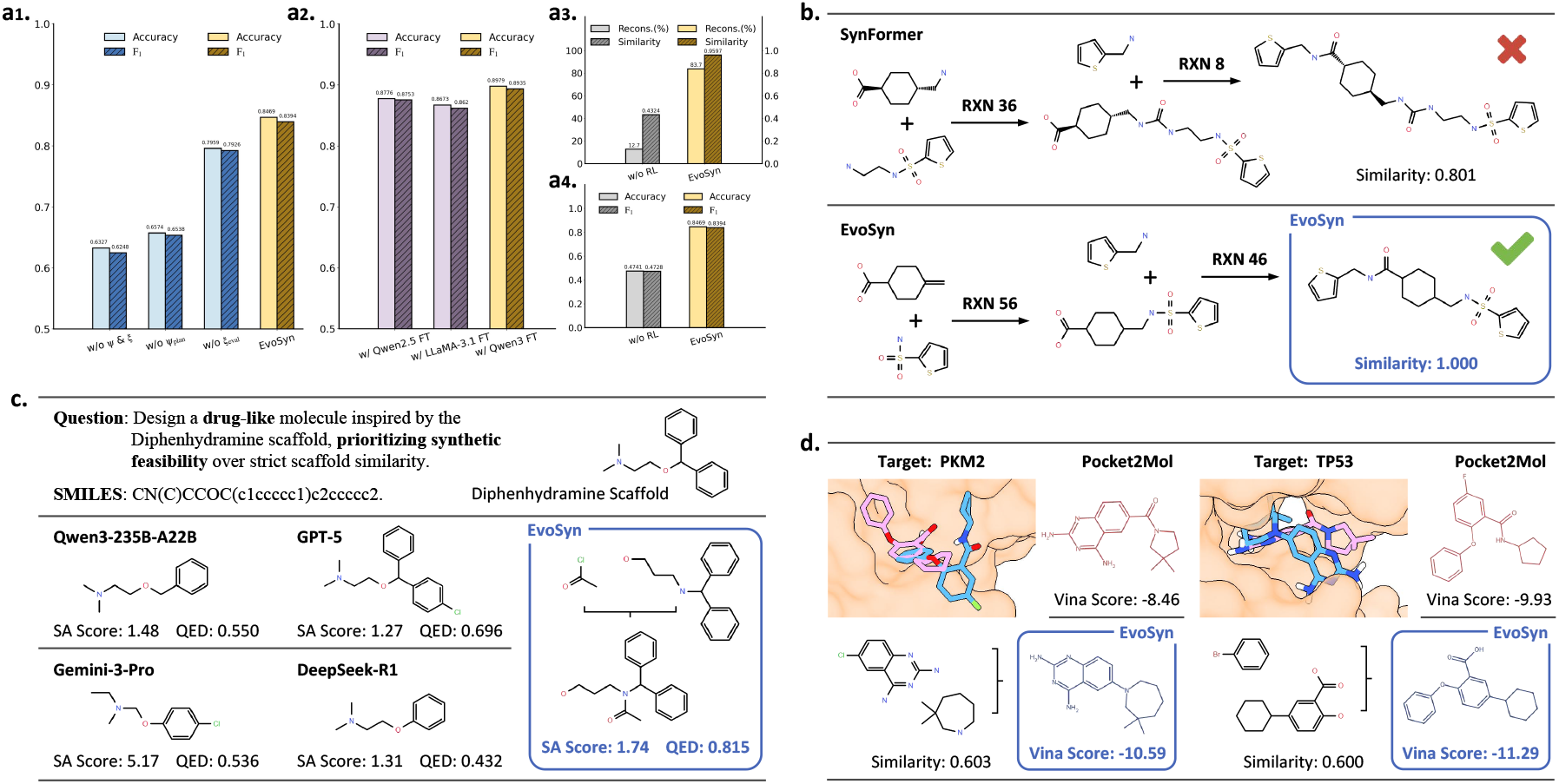
**a1-a4**.Ablation Study of EvoSyn. **a1**.The contribution of the co-evolving agents. **a2**.The impact of self-evolving with fine-tuning. **a3-a4**.The contribution of self-evolving with reinforcement learning. **b**.EvoSyn outperforms baseline in multi-step planning. **c**.Preference-aware synthesis of EvoSyn and LLMs. **d**.Visualization of synthesizable lead optimization.

Complementing the planner, the Consistency Evaluator *ξ*_eval_ plays an indispensable role in the refinement stage. The performance gap observed in EvoSyn (w/o *ξ*_eval_) verifies that relying solely on one-shot generation is insufficient, and iterative feedback is requisite for mitigating hallucinations and correcting logical inconsistencies. Furthermore, the minimal performance of the variant EvoSyn (w/o *ψ*_plan_ & *ξ*_eval_), which lacks both agents, confirms that the superior capability of EvoSyn stems not from individual components alone, but from the synergistic co-evolution of global planning and dynamic evaluation.

#### Impact of self-evolving with fine-tuning

We further assess the scalability of EvoSyn by comparing its zero-shot performance against versions fine-tuned (FT) via our self-evolving strategy. Remarkably, the zero-shot EvoSyn already achieves a high Accuracy of 0.8469, demonstrating the robustness of the co-evolution. When equipped with fine-tuning, the performance is significantly elevated. As shown in Figure 2 a2 and Supplementary Table S3, the LLaMA-3.1-8B based variant reaches 0.8673, while the Qwen2.5-7B based variant peaks at 0.8776. Consequently, our final EvoSyn model with Qwen3-4B achieves a state-of-the-art Accuracy of 0.8979.

These results indicate a complementary relationship: while the co-evolution mechanism serves as a powerful reasoning engine during inference, the self-evolution mechanism enables the model to permanently internalize domain-specific patterns, pushing the performance to the optimal level.

#### Contribution of self-evolving with reinforcement learning

To investigate the contribution of the RL component, we isolate it and denote the variant as EvoSyn (w/o RL). As shown in Figure 2 a3 and Supplementary Table S4, on the Enamine REAL subset of the Molecule Reconstruction task, EvoSyn (w/o RL) exhibits a significant performance drop due to the lack of policy guidance. In Figure 2 a4, without the RL component, the Synthesis Planner *ψ*_plan_ fails to generate correct synthesis pathways and thus cannot properly support the Reasoning Responder *ρ*_res_, leading to further performance degradation in SynQA-Basics.

Therefore, relying solely on the co-evolution of agents is insufficient to achieve preference-aware molecular synthesis. The molecular synthesis algorithm we propose, based on a Markov Game formulation, provides essential support for agent reasoning.

### 4.6. Case study

To intuitively understand how EvoSyn navigates complex chemical spaces, we present three qualitative case studies visualizing its performance in multi-step planning, preference-aware synthesis, and lead optimization.

#### Multi-step planning

Figure 2 b and Supplementary Figure S1 present the multi-step synthesis planning of EvoSyn and SynFormer for the same molecule. Benefiting from the Markov Game for synthesis, EvoSyn accurately planned the correct synthetic route. In contrast, since SynFormer requires predicting products from scratch, slight deviations at the beginning result in significant divergence in the subsequent synthesis direction.

#### Preference-aware synthesis

Figure 2 c illustrates a query prioritizing *drug-likeness* and *synthetic feasibility* over strict scaffold similarity. Generalist LLMs like Gemini-3-Pro fail this constraint, generating hard-to-synthesize molecules (SA score 5.17) without synthesis pathways. In contrast, EvoSyn generates a molecule with a superior QED score (0.815 vs. 0.696 for GPT-5) while maintaining a low SA score (1.74). Crucially, by explicitly outputting the precursor reactants (blue box), EvoSyn verifies synthesizability by design, proving its capability to generate actionable chemical plans rather than mere theoretical structures.

#### Synthesizable lead optimization

Figure 2 d visualizes the optimization of Pocket2Mol lead candidates. 3D superposition confirms that the molecules optimized by EvoSyn (blue) achieve deeper binding pocket penetration than the reference leads (pink), corresponding to significant bioactivity gains (Vina Score improves from -9.93 to -11.29). Unlike baselines that may compromise structural integrity for energy minimization, EvoSyn maintains high structural similarity (∼0.600) while providing exact reaction precursors. This ensures the optimized candidates are not just “virtual hits” with low energy, but synthetically feasible leads grounded in chemical reality.

## 5. Conclusion

We presented EvoSyn, an evolutionary multi-agent framework that bridges linguistic reasoning and chemical validation through a dual-process evolutionary paradigm. By integrating collaborative constraint alignment with game-theoretic self-evolution, EvoSyn achieves state-of-the-art performance, generating molecules that are both bioactive and synthetically actionable. Future work will extend the framework to complex multi-step synthesis and real-world experimental settings by reducing current reliance on computational rewards.

## Supplementary data

Supplementary data is available at Bioinformatics online.

## Data availability

The data and implementation code during this study are included in the supplementary information files.

## Acknowledgments

The authors thank the anonymous reviewers for their valuable suggestions. This study has been supported by the National Natural Science Foundation of China [T2394502], the Guangdong S&T Program [2023B1111030002, 2024B1111140001], the Shenzhen Science and Technology Plan Project [CJGJZD20220517142201004], the National Natural Science Foundation of China [62302537, 625007225], the Guangdong Basic and Applied Basic Research Foundation [2025A1515060011], the Guangzhou Basic and Applied Basic Research Foundation [2024A04J4449], and the Lingang Laboratory [LGL-8888].

